# A*t*FZL is required for correct starch granule morphology in Arabidopsis chloroplasts

**DOI:** 10.1101/2022.10.20.512996

**Authors:** Lara Esch, Qi Yang Ngai, J. Elaine Barclay, David Seung

## Abstract

The control of starch granule number and morphology in plastids is poorly understood. Here, we demonstrate that *At*FZL, a protein involved in thylakoid membrane organisation, is required for correct starch granule morphology in Arabidopsis. Leaves of mutants lacking *At*FZL had the same starch content as wild-type leaves, but their starch granules were smaller and had a distinct, uneven surface morphology. Most chloroplasts in the mutant were larger than those of the wild type.

However, the difference in chloroplast size could not explain the difference in granule size and shape in the *Atfzl* mutants, since other mutants with larger chloroplasts than the wild type (*arc* mutants) had granules that were similar in size and shape to wild-type granules. As reported previously, the *Atfzl* mutant had aberrant thylakoid organisation. We found that this phenomenon was particularly pronounced in regions surrounding starch granules. The location of the thylakoid-bound granule initiation protein MFP1 was unaffected in the *Atfzl* mutant. We propose that *At*FZL affects starch granule size and shape by influencing thylakoid organisation at the periphery of starch granules. Our results are consistent with an important role for thylakoid architecture in determining granule morphology.

## Introduction

Starch is the primary form of carbohydrate storage in plants. Plants use photoassimilates to synthesise semi-crystalline insoluble starch granules within plastids of leaves and storage organs. In leaves, starch granules are synthesized in chloroplasts during the day then degraded during the night to supply metabolic energy required for maintenance and growth in the absence of photosynthesis (Smith and Zeeman, 2020). In most plants, leaf starch granules are relatively small and have a flattened lenticular shape. In Arabidopsis, each chloroplast is reported to contain 5-7 starch granules that are approx. 1-2 µm in diameter at the end of the day (Crumpton-Taylor *et al*., 2012) and are formed within stromal pockets between thylakoid membranes (Seung *et al*., 2018; Bürgy *et al*., 2021). However, factors involved in determining the number, shape and size of starch granules within leaf chloroplasts are still poorly understood.

Recent work has advanced our understanding of starch granule initiation in chloroplasts, which determines both the number and size of granules. In Arabidopsis, the key enzyme that mediates granule initiation is STARCH SYNTHASE 4 (*At*SS4) – a glucosyltransferase that may perform the initial glucan elongation steps that prime granule formation (Roldán *et al*., 2007). Mutants defective in *At*SS4 have drastic reductions in the number of starch granules in chloroplasts, most containing zero or one granule (Roldán *et al*., 2007; Bürgy *et al*., 2021; Lu *et al*., 2018). Several additional proteins are proposed to act in granule initiation, by influencing *At*SS4 activity and/or localisation. These include PROTEIN TARGETING TO STARCH2 (*At*PTST2), a Carbohydrate Binding Module 48 (CBM48)-containing protein proposed to deliver short maltooligosaccharide (MOS) substrates to *At*SS4 for further elongation (Seung *et al*., 2017). *At*PTST2 interacts with the MAR1-BINDING FILAMENT PROTEIN (*At*MFP1) (Seung *et al*., 2018). Most chloroplasts in mutants defective in *At*PTST2 or *At*MFP1 have a single, large granule, but the granules retain a normal flattened disc shape (Seung *et al*., 2018; Thieme *et al*., 2022). Interestingly, in *Atss4* mutants, starch granules are round rather than lenticular, suggesting AtSS4 is also required for correct granule morphology (Roldán *et al*., 2007; Bürgy *et al*., 2021; Lu *et al*., 2018).

It has also been speculated that the flattened shape of leaf starch granules may be influenced by space available for granule growth within the chloroplast, in between layers of thylakoid membranes (Zeeman *et al*., 2002). Indeed, recent work demonstrated that granules form within stromal pockets between thylakoids, in which they initiate as multiple small granules that eventually fuse into a single granule (Bürgy *et al*., 2021). There is also evidence for associations between granule initiation proteins and thylakoid membranes. *At*MFP1 is exclusively bound to the stromal side of thylakoid membranes (Jeong *et al*., 2003; Seung *et al*., 2018). This allows some *At*PTST2 to attach to thylakoids through interaction with *At*MFP1 (Seung *et al*., 2018). *At*MFP1 and *At*PTST2 co-locate to numerous, discrete puncta within chloroplasts, and it is hypothesised that they define areas in which granules initiate and form (Seung *et al*., 2017; Seung *et al*., 2018; Seung and Smith, 2019). *At*MFP1 may also associate with plastid nucleoids (Jeong *et al*., 2003), although the role of this interaction is not known.

These previous findings raise the possibility that the number and shape of starch granules are influenced by the nature and extent of stromal pockets that form between thylakoids. Relatively little is known about the nature of these pockets and the factors involved in their formation, but important clues come from the phenotypes of Arabidopsis mutants deficient in a chloroplastic dynamin-like protein, FZO-LIKE (FZL). *At*FZL is located in punctate structures associated with both thylakoids and envelope membranes of chloroplasts. In mutants lacking this protein, chloroplast thylakoids are disorganised and the ratio of granal to stromal thylakoids is reduced (Gao et al., 2006). It has been proposed that *At*FZL may fuse membrane compartments together during thylakoid biogenesis, with a particular role in fusing developing grana stacks to stromal thylakoids (Gao et al. 2006; Liang et al., 2018). Other studies of Arabidopsis and *Chlamydomonas fzl* mutants have confirmed the importance of FZL for chloroplast structure (Findinier *et al*., 2019; Landoni *et al*., 2013).

Here, we have taken advantage of the *Atfzl* mutant to analyse whether disruption of thylakoid organisation affects starch granule formation and morphology. We demonstrate that *At*FZL is required for correct starch granule morphology in Arabidopsis. Chloroplasts of the *Atfzl* mutant produce starch granules that are reduced in size and have uneven surface morphology. We propose that these changes in granule morphology result from disorganised thylakoids at the periphery of starch granules in the mutant, indicating that thylakoid architecture influences starch granule growth.

## Results

### Loss of AtFZL results in increased starch granule number per chloroplast due to increased chloroplast size

To investigate whether the thylakoid-organising protein *At*FZL influences starch granule formation in Arabidopsis leaves, we obtained three independent T-DNA mutants from the Eurasian Arabidopsis Stock Centre (uNASC). Two of these lines have been described previously (Gao *et al*., 2006): *Atfzl-1* (SALK_009051) has two insertions in the *AtFZL* gene (one in the intron between exons 9 and 10 that encodes the second transmembrane domain, and another in the 3’ UTR), while *Atfzl-2* (SALK_033745) has an insertion in exon 2 (Fig. 1a). An additional line, *Atfzl-3* (SALK_152584C) has a T-DNA insertion in exon 4, which encodes part of the GTP-binding domain. The presence of each T-DNA insertion in the respective mutants was confirmed using PCR. All plants were grown in controlled environment chambers under a 12 h photoperiod.

**Figure 1:**
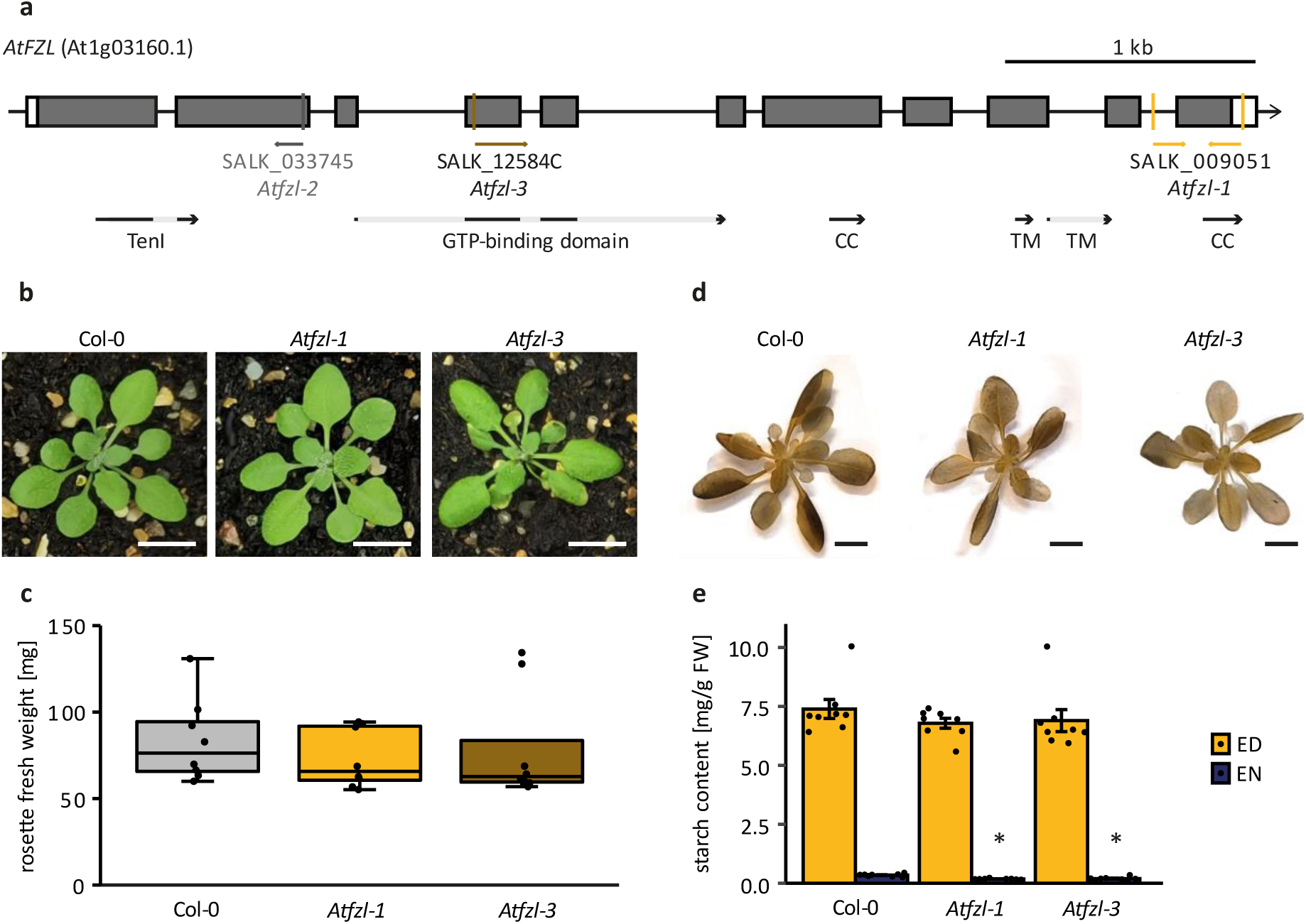
Starch content of Arabidopsis *fzl* mutants. **(a)** Schematic illustration of the gene model for the primary transcript of *AtFZL* (At1g03160.1). Exons are represented by grey boxes and UTRs are represented by white boxes. Grey and yellow arrows indicate T-DNA insertion sites (SALK_009051: *Atfzl-1*, SALK_033745: *Atfzl-2*, SALK_12584C: *Atfzl-3*). Regions encoding protein domains are shown as black and grey arrows (CC: coiled coil, TenI: thiamine phosphate synthase, TM: transmembrane). **(b)** Photographs of 30-day-old *Atfzl* mutant and wild-type (Col) rosettes. Bar = 1 cm. **(c)** Fresh weight of 28-day-old rosettes. Dots represent fresh weight of an individual rosette (*n* = 8 per genotype). The bottom and top of the box represent the lower and upper quartiles respectively, and the band inside the box represents the median. The ends of the whiskers represent values within 1.5x of the interquartile range. Outliers are values outside 1.5x the interquartile range. No significant differences (including outliers) were observed under Kruskal-Wallis One-Way Analysis of Variance on the Ranks (P = 0.437). **(d)** Iodine-stained, 33-day-old *Atfzl* mutant and wild-type rosettes (Col-0). Bar = 1 cm. **(e)** Starch content of *Atfzl* mutants at the end of the day (ED) and the end of the night (EN). Bars and error bars represent the mean±SEM of *n* = 8 plants, while dots represent individual data points. Values indicated with an asterisk are significantly different from Col-0 at each timepoint under a one-way ANOVA and Holm-Šídák multiple comparisons procedure (P ≤ 0.001).

Although previous studies reported pale leaves, chlorotic lesions and delayed flowering in *Atfzl* mutants (Gao et al., 2006; Landoni et al., 2013), rosette morphology of the *Atfzl-1* and *Atfzl-3* mutants was indistinguishable from the wild type (WT) under our growth conditions (Fig. 1b).

The fresh weights of the *Atfzl-1* and *Atfzl-3* rosettes were not significantly different from those of the WT (Fig. 1c). We then assessed the effect of the *Atfzl* mutations on transient starch accumulation in leaves. Iodine staining for starch in whole rosettes harvested at the end of the day showed no obvious differences in staining pattern or intensity between the *Atfzl* mutants and the WT (Fig. 1d). Quantitative starch assays confirmed that there were no differences in starch content between the mutants and the WT at the end of the day (Fig. 1e). However, there was a significant reduction in the end-of-night starch content in both mutants compared to the WT.

To determine if *Atfzl* mutations affect the number and size of starch granules per chloroplast, we examined sections of rosette leaves using light microscopy. Consistent with earlier reports (Gao *et al*., 2006), chloroplasts in mesophyll cells in the *Atfzl* mutants were larger than those of the WT (Fig. 2a-d). The total chloroplast area per cell section was significantly larger in the *Atfzl-3* mutant and slightly larger in the *Atfzl-1* mutant, when compared to the WT (Fig. 2a). Additionally, the total area of individual mesophyll cells was significantly larger in the *Atfzl-3* mutant and slightly larger in the *Atfzl-1* mutant than in the WT (Fig. S2d). Moreover, the shape of the chloroplasts in the *Atfzl* mutants was drastically altered and not uniform - many were flat and elongated, while some were WT-like and round (Fig. 2b-d).

**Figure 2:**
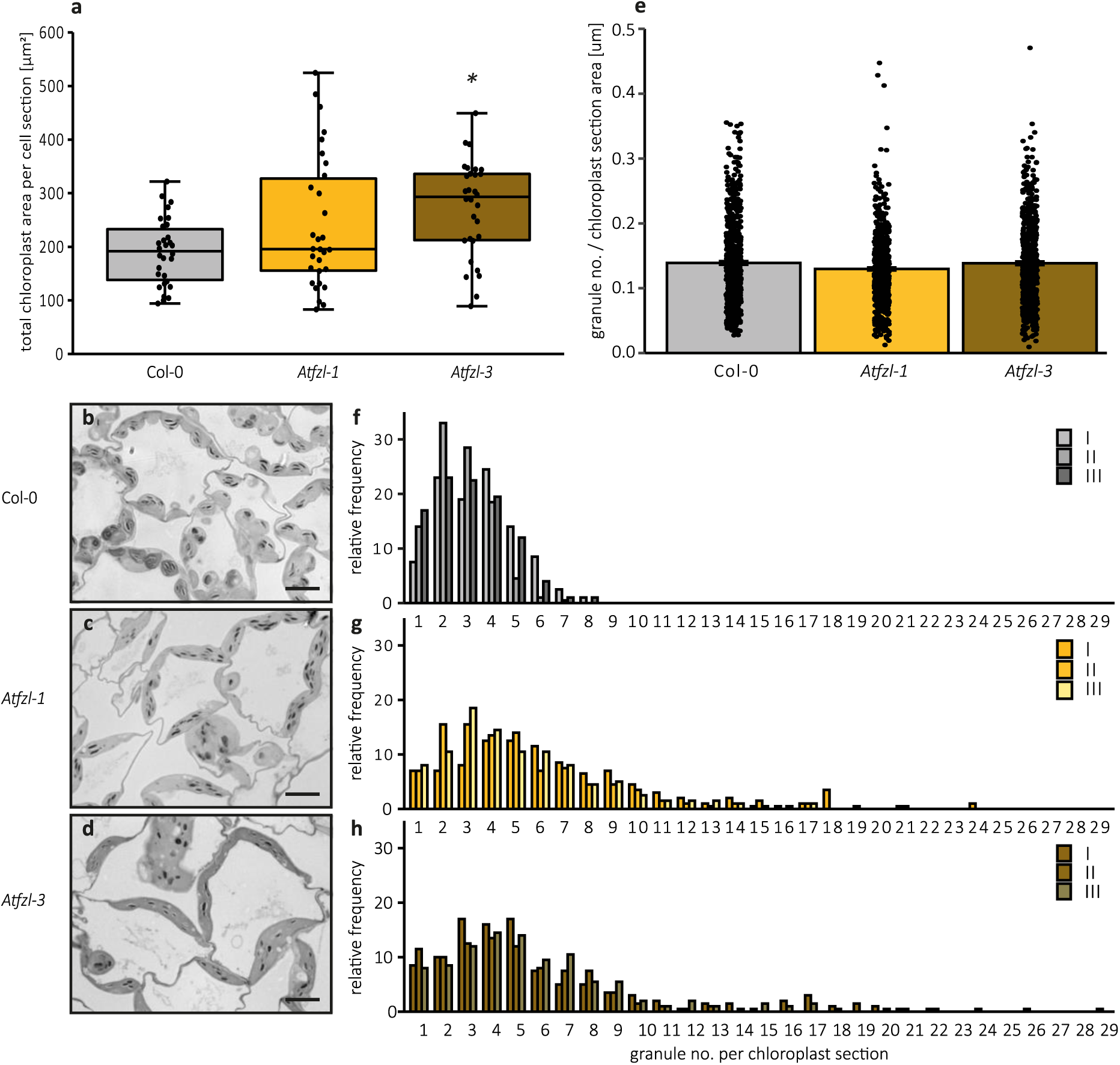
Starch granule number per chloroplast section in *Atfzl* mutants. **(a)** Total chloroplast section area per cell section. Values represent total area of chloroplast sections per cell for ten cells per biological replicate. Values that are significantly different from Col-0 under a Kruskal-Wallis One Way Analysis of Variance on Ranks followed by a post-hoc ukey’s est (P ≤ 0.0) are indicated with an asterisk. **(b-d)** Light microscopy images of leaf sections of *Atfzl* mutants and the wild type (Col-0). A recently fully expanded leaf was harvested from 29-day-old plants at the end of day. Sections were stained ith toluidine lue and chi ‘s stain. Bar = 10 μm. **(e)** Number of granules expressed relative to the chloroplast section area. No significant differences between genotypes were observed under Kruskal-Wallis One-Way Analysis of Variance on the Ranks (P = 0.140). **(f-h)** Starch granule number per chloroplast section. Histograms show the frequency of chloroplasts containing a given number of granule sections relative to the total number of chloroplasts analysed (200 chloroplasts per replicate). Three biological replicates were analysed per genotype (labelled I-III), where each was prepared from a different plant.

Chloroplasts in the *Atfzl* mutants had visibly more starch granules than those of the WT. We therefore quantified the number of starch granules per chloroplast section using light microscopy images. The number of starch granules, when expressed relative to the chloroplast section area, was identical between the *Atfzl* mutants and the WT (Fig. 2e). Therefore, given the heterogeneity in chloroplast size observed in the mutant, the distribution of granule numbers per chloroplast section for the *Atfzl-1* and *Atfzl-3* mutants was significantly different from that of the WT – with the mutants having a broader distribution that was shifted towards higher granule numbers (Fig. 2f-h). We fitted a negative binomial regression to these distributions and extracted the mean count of granules per chloroplast section. *Atfzl-1* and *Atfzl-3* had a predicted mean count of 5.69 and 5.68 granules per chloroplast section, respectively; compared to the WT which had a predicted mean of 3.10 granules per chloroplast section. A similar trend towards increased numbers of granules per chloroplast was observed for the *Atfzl-2* mutant (Fig. S1). When we correlated the number of starch granules per chloroplast section to the section area of the chloroplasts, we observed a distinct tendency of larger chloroplasts containing more starch granules in the *Atfzl* mutants (Fig. S2a-c), suggesting that chloroplast size was the primary factor underpinning the differences in the number of granules observed per chloroplast in the mutants.

### AtFZL is required for normal starch granule morphology

We then examined starch granule morphology in the *Atfzl-1* and *Atfzl-3* mutants. Leaf starch was purified from whole rosettes harvested at the end of the day. Analysis of starch granule size distribution on the Coulter counter showed that *Atfzl-1* and *Atfzl-3* mutants had smaller granules than the WT (Fig. 3a). In the *Atfzl-1* and *Atfzl-3* mutants, the average granule diameter was 1.765±0.032 µm and 1.817±0.004 µm respectively. In the WT, the mean granule diameter was 2.120±0.024 µm. The smaller granule size in the mutants was also evident when purified starch granules were observed using scanning electron microscopy (Fig. 3 b-h). Interestingly, starch granules of the *Atfzl* mutants had distinctive uneven surfaces, as opposed to the smooth surfaces of WT granules. This feature was more pronounced in granules of *Atfzl-3* than in those of *Atfzl-1*.

**Figure 3:**
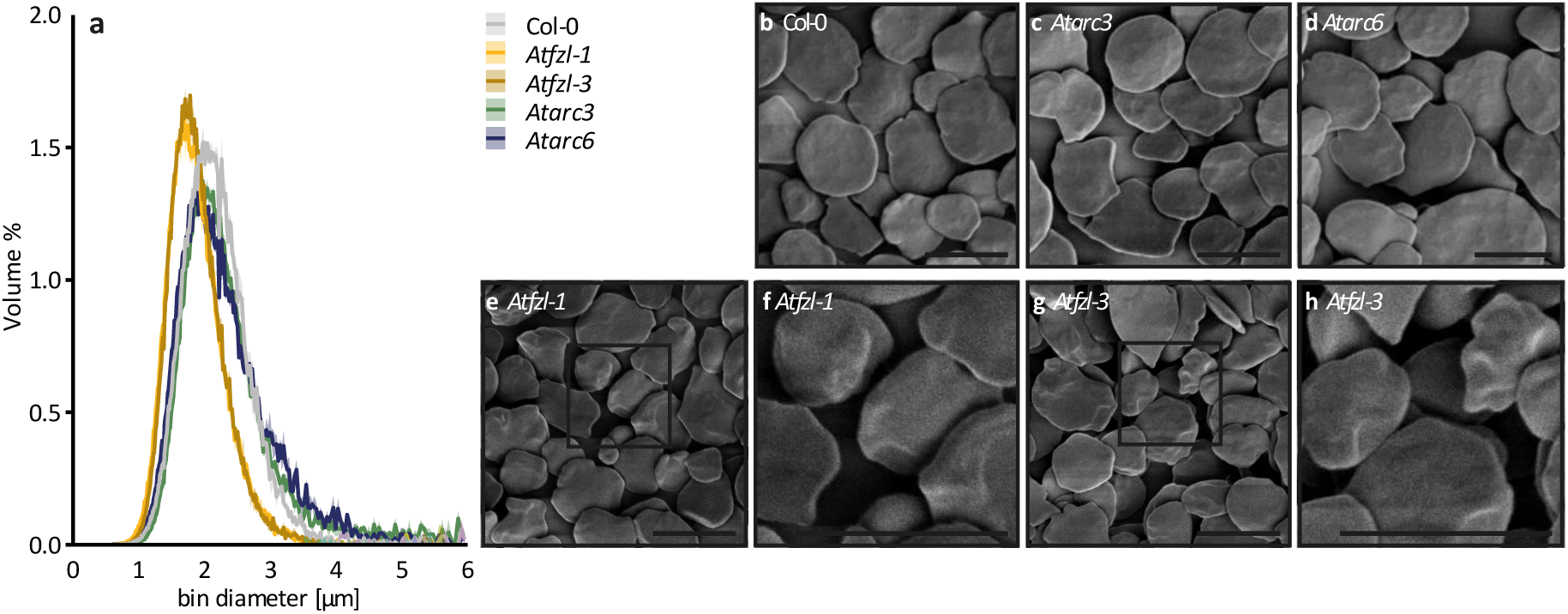
Starch granule morphology of *Atfzl* mutants. **(a)** Size distributions of starch granules purified from 34-day-old rosettes harvested at the end of the day, pooling 60 rosettes per genotype. The volume of granules at each diameter relative to the total granule volume was quantified using a Coulter counter. Values represent mean (solid line) ± SEM (shading) of three technical replicates. **(b-h)** SEM images of starch granules purified from 34-day-old rosettes. Bar = μm. Panels (f) and (n) are close-up images of *Atfzl-1* and *Atfzl-3*, showing the uneven granule morphology.

To determine whether the altered starch granule size and morphology in the *Atfzl* mutants was an inevitable consequence of larger chloroplast size in the mutants, we also examined purified starch granules from *Atarc3* and *Atarc6* mutants. Both these mutants are defective in plastid division and have larger chloroplasts (chloroplast cross-sectional areas are about two and four times larger than WT in *Atarc3* and *Atarc6*, respectively), and like our *Atfzl* mutants, the number of granules per chloroplast volume in the *Atarc3* and *Atarc6* mutants is identical to the WT (Crumpton-Taylor *et al*., 2012).

However, the arrangement of the thylakoid membranes in *Atarc3* and *Atarc6* mutants remains largely unaltered (Maple *et al*., 2007; Pyke *et al*., 1994; Robertson *et al*., 1995; Shimada *et al*., 2004; Vitha *et al*., 2003). Coulter counter analysis of the purified starch granules of the *Atarc3* and *Atarc6* mutant showed that their sizes were not statistically different from the WT, with mean granule diameters of 2.349±0.031 and 2.149±0.010 µm, respectively (Fig. 3a). *Atarc3* and *Atarc6* starch granules also had similar morphology to WT granules (Fig. 3b). This indicates that the altered size and shape of starch granules in *Atfzl* mutants is not a function of increased chloroplast size, but may be attributable to altered thylakoid structure and organisation in these mutants.

### The Atfzl-2 and Atfzl-3 mutants have altered thylakoid organisation around starch granules

We next examined the organisation of stromal pockets and starch granules within chloroplasts of the *Atfzl* mutants. Consistent with previous reports (Gao *et al*., 2006), transmission electron microscopy (TEM) showed disorganised thylakoids in chloroplasts of the *Atfzl* mutants. Granal lamellae appeared less stacked and uniform, and spaces between the thylakoid membranes were larger and more randomly distributed. Starch granules appeared to be less evenly distributed throughout the chloroplast in the *Atfzl* mutants, with some clusters of granules close to each other and some granules occurring unusually close to the plastid envelope. Most strikingly, membrane disorganisation was particularly pronounced around starch granules, often appearing as detached membrane structures, likely thylakoid membranes, in unusually large pockets surrounding the starch granules in the *Atfzl-3* mutant (Fig. 4). These thylakoid membrane phenotypes were also seen in the *Atfzl-2* mutant (Fig. S3), but not in the *Atfzl-1* mutant, possibly reflecting different degrees of phenotype severity depending on the location of the T-DNA insertion site within the *FZL* gene (Fig. 1a).

**Figure 4:**
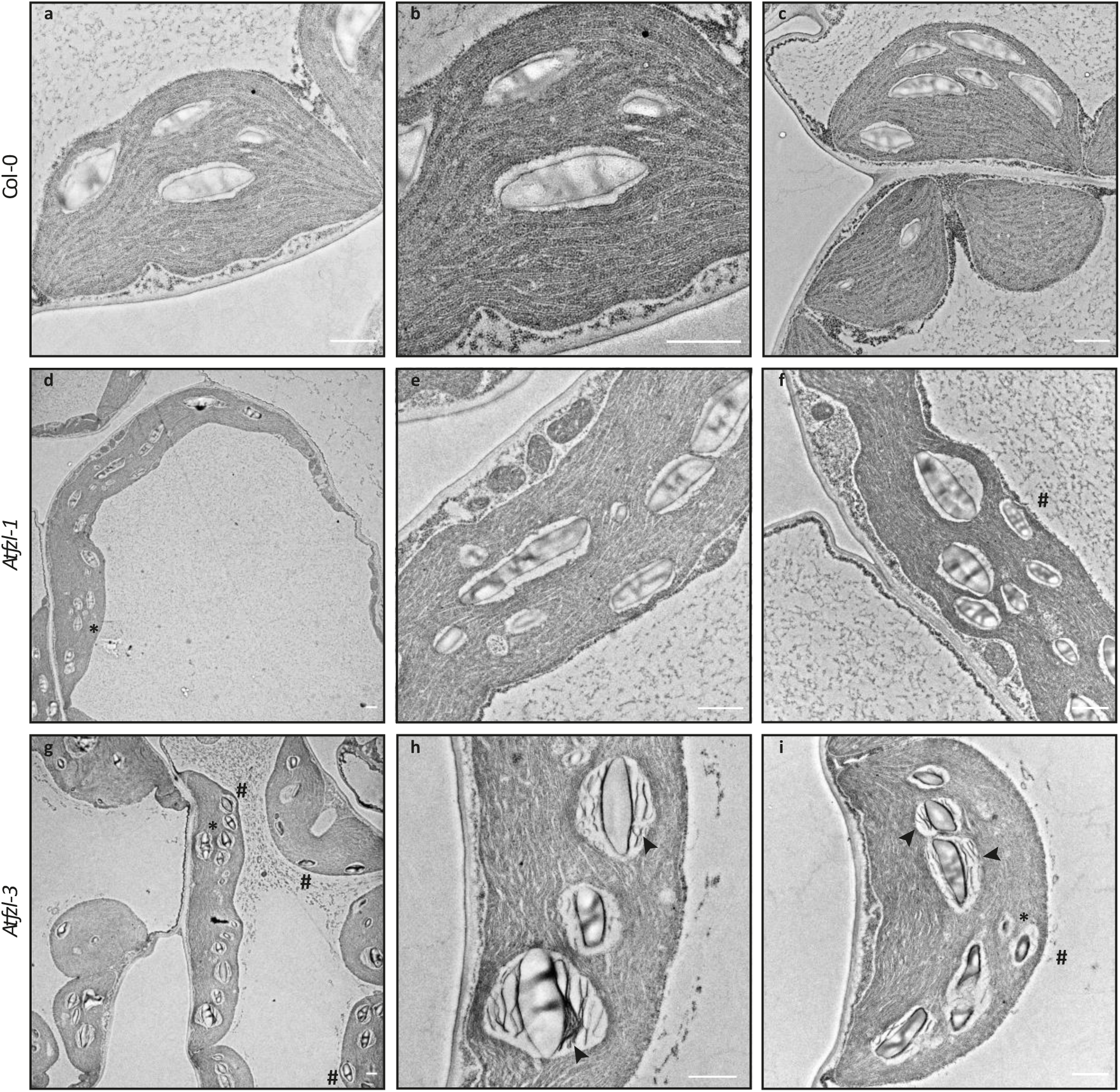
Chloroplast ultrastructure and starch granule placement in *Atfzl* leaves. TEM images of thin sections prepared from recently fully expanded leaves harvested from 29-day-old rosettes at the end of the day. Arrowheads indicate disor anised, ‘detached’ thylakoid layers observed at the periphery of starch granules in *Atfzl-3*. Asterisk (*) indicates granule clusters. Hash (#) indicates granules close to chloroplast envelope. Bar = 1 μm

Given the unusual membrane organisation around starch granules, we also examined thylakoid organisation and plastid ultrastructure in leaves of the *Atfzl* mutants harvested at the end of the night, after starch had been largely depleted (Fig. 5). Starch granules were almost completely degraded in the *Atfzl* mutants and the pockets previously occupied by granules were greatly reduced in size, but the detached thylakoid membranes observed around granules in *Atfzl-3* chloroplasts now occupied the stromal pocket. By contrast, in the WT, the thylakoid pockets around starch granules had shrunk to accommodate the reduced granule size but the thylakoids remained organised.

**Figure 5:**
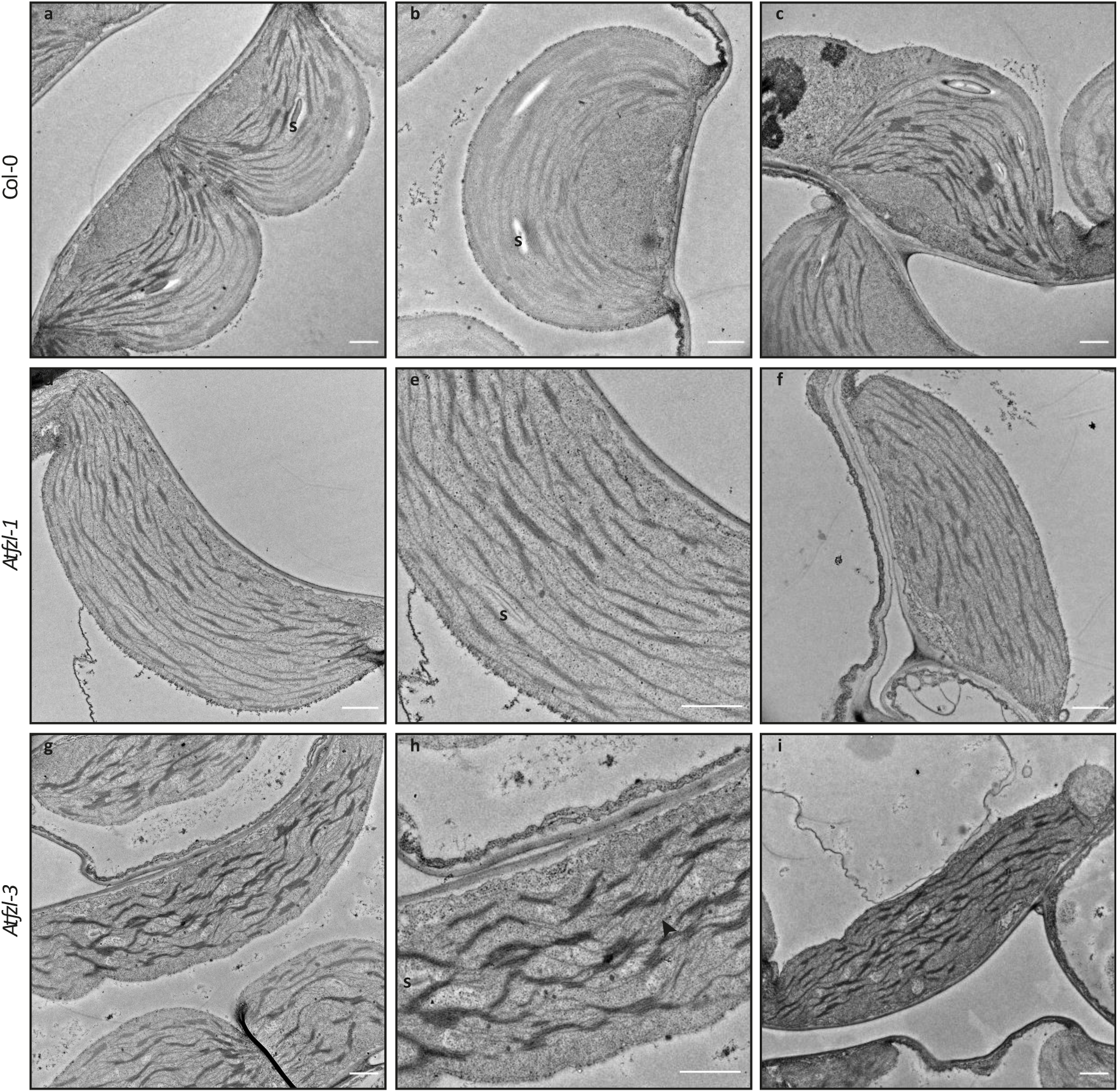
Chloroplast ultrastructure after starch granule degradation. TEM images of thin sections prepared from recently fully expanded leaves harvested from 32-day-old rosettes at the end o the ni ht. rro heads indicate disor anised, ‘detached’ thylakoid layers running through stromal pockets. “s” indicates starch ranules. Bar = 1 μm

### The punctate localisation of AtMFP1 is not dependent on AtFZL

Since thylakoid organisation was severely disrupted in the *Atfzl* mutants, we investigated whether there is functional link between *At*FZL and MAR BINDING FILAMENT-LIKE PROTEIN 1 (MFP1), a thylakoid-associated protein that is important for starch granule initiation in Arabidopsis.

Localisation experiments with *At*FZL transiently expressed in *Nicotiana benthamiana* with a C-terminal YFP tag (*At*FZL-YFP) showed that it localises to small puncta in chloroplasts (Fig. 6a-c), as has been reported previously (Gao *et al*., 2006). *At*MFP1 also localises to discrete puncta (Seung *et al*., 2018), but co-expression of *At*FZL-YFP and *At*MFP1-RFP did not show a clear co-localisation pattern between the punctae of the two proteins (Fig. 6d-h). A pairwise immunoprecipitation assay between the *At*FZL-YFP and *At*MFP1-RFP proteins co-expressed in *N. benthamiana* leaves also failed to reveal an interaction between the two proteins (Fig. S4). We then tested whether *At*FZL is necessary for the correct localisation of *At*MFP1 in Arabidopsis. We created transgenic Arabidopsis lines expressing *At*MFP1-YFP in the WT and *Atfzl-3* mutant background. *At*MFP1-YFP formed discrete puncta in both mesophyll and epidermal chloroplasts of the mutant as well as the WT. Interestingly, although it was visible from the chloroplast autofluorescence that the *Atfzl-3* mutant had drastically increased chloroplast size in the mesophyll cells compared to the WT, the chloroplasts in the epidermal pavement cells were indistinguishable in size and shape from those of the WT (Fig. 6i-n). Taken together, it is unlikely that *At*FZL affects starch granule morphology by affecting *At*MFP1 function.

**Figure 6:**
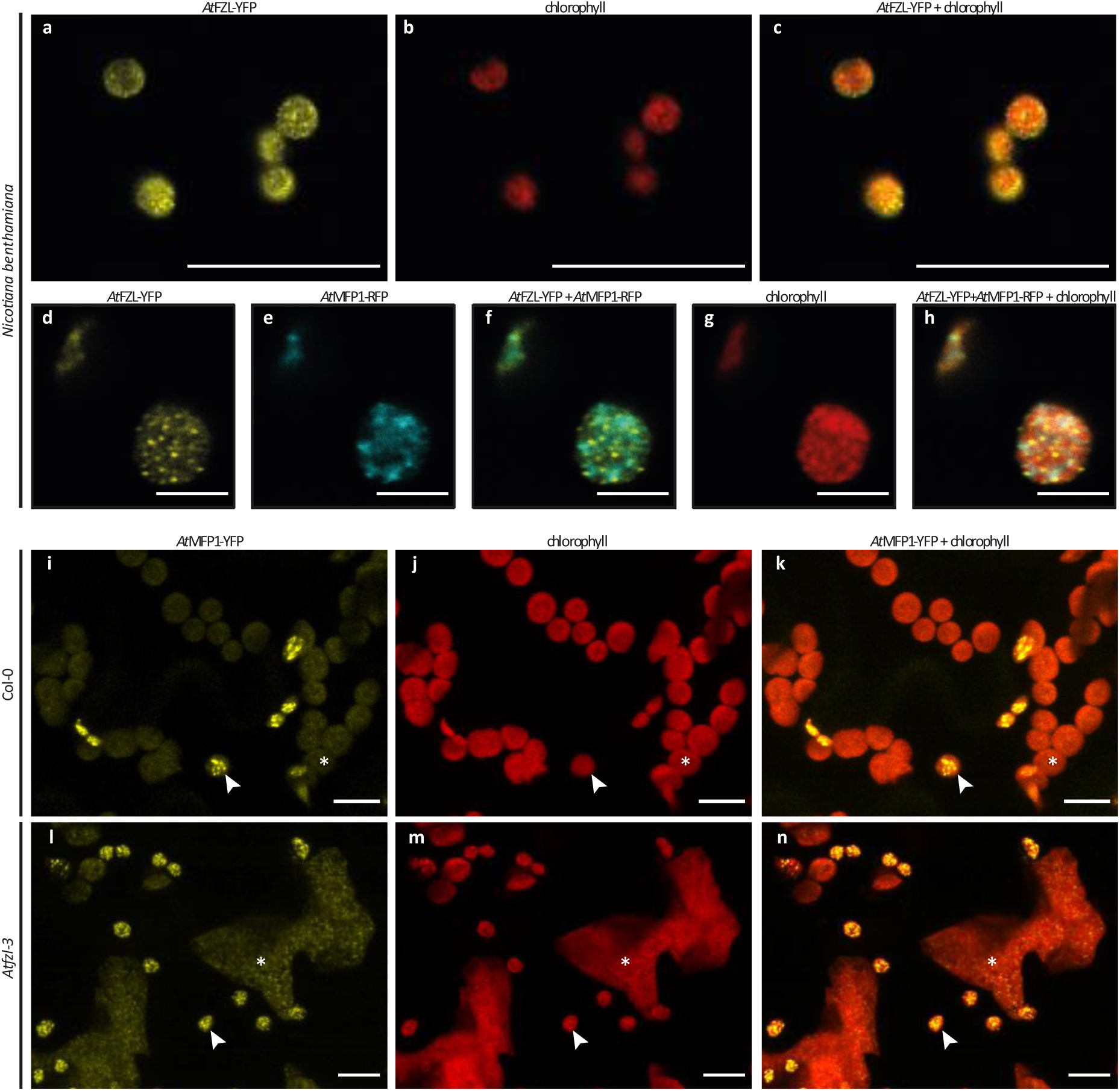
Localisation of *At*FZL and *At*MFP1. Images were obtained using confocal laser-scanning microscopy. **(a-h)** Images of *At*FZL-YFP and *At*MFP1-RFP expressed in *N. benthamiana* epidermal cells. The YFP- and RFP fluorescence are shown in yellow and cyan, while chlorophyll autofluorescence is shown in red. **(i-n)** Z-projection of image stacks of *At*MFP1-YFP stably expressed in Arabidopsis wild-type (Col-0) and *Atfzl-3* backgrounds. Recently fully expanded leaves from 5-week-old rosettes were imaged. Examples of chloroplasts of epidermal pavement cells are indicated with an arrowhead, and those of mesophyll cells are marked with an asterisk. Bars = 5 μm.

## Discussion

### Correct thylakoid organisation is required for normal starch granule morphology in chloroplasts

We demonstrate that *At*FZL, a chloroplast protein required for correct organization of thylakoid membranes (Gao et al., 2006; Liang et al., 2018), is also necessary for correct starch granule morphology. Starch granules in the *Atfzl* mutants are smaller and have less even surfaces than those of the wild type (WT)(Fig. 3).

Starch granules in Arabidopsis chloroplasts orm ithin de ined ‘stromal pockets’ et een thylakoid membranes, via the fusion of multiple, separately-initiated starch granules (Bürgy *et al*., 2021). The alterations in thylakoid structure in *Atfzl* mutants appear to affect the size and structure of these stromal pockets. Consistent with previous reports (Gao *et al*., 2006), we found that *Atfzl* chloroplasts had disorganised thylakoids with less uniformly stacked grana than WT chloroplasts, and more randomly distributed intermembrane spaces (Fig. 4). In the *Atfzl-2* and *Atfzl-3* mutants, we also observed disorganised or detached membranes, likely thylakoid membranes, particularly within the stromal pockets around the starch granules (Fig. 4). These membranes were still visible within stromal pockets at the end of the night following starch degradation (Fig. 5). Interestingly, detached membranes were not observed in the *Atfzl-1* mutant (Fig. 4 and Fig. S3). In *Atfzl-1*, the T-DNA insertion is integrated in the region encoding the C-terminal end of *At*FZL after the predicted transmembrane domains. Insertions in both *Atfzl-2* and *Atfzl-3* are closer to the start of the coding sequence, either before or within the region encoding the GTP-binding domain (Fig. 1). It is plausible that the position of the *Atfzl-1* insertion towards the end of the coding sequence permits the translation of a partly functional protein, resulting in a weaker phenotype. Notably, the Arabidopsis gene annotation for *FZL* suggests four different splice variants at the locus, two of which (At1g03160.2 and At1g03160.3) encode a truncated protein lacking the two C-terminal transmembrane domains. The *fzl-1* mutation does not disrupt the coding sequence of these two splice variants, and may allow these truncated proteins to be expressed.

We propose that the aberrant starch granule size and shape observed in the *Atfzl* mutants are consequences of altered thylakoid structure. Disrupted structure of the stromal pockets may limit either the space available for fused granules within the pocket to grow, or the number of initiation events that can occur within a stromal compartment – resulting in smaller starch granules. The distinct uneven surface of the mutant starch granules, which was more pronounced in the *Atfzl-3* mutant than in *Atfzl-1*, might also be influenced by the less organised thylakoid membranes around the stromal pocket, which may interfere with starch granule growth and prevent the formation of a smooth granule surface. It is also possible that restricted stromal space around growing starch granules might introduce tension within the matrix of growing granules, leading to deformation.

Starch granule morphology is in addition influenced by components that control the non-random deposition of new material (Bürgy *et al*., 2021). Arabidopsis STARCH SYNTHASE 4 (*At*SS4), a key enzyme in starch granule initiation (Roldán *et al*., 2007), is required for anisotropic growth of starch granules in Arabidopsis (Bürgy *et al*., 2021). Knock-out of *At*SS4 results in uniform deposition of newly synthesised starch onto the granule surface, leading to spherical granules in place of the lenticular granules in the WT (Bürgy *et al*., 2021; Roldán *et al*., 2007). *At*SS4 and PROTEIN TARGETING TO STARCH2 (*At*PTST2) are associated with thylakoid membranes (Gámez-Arjona *et al*., 2014). *At*PTST2 co-localises in discrete puncta within Arabidopsis chloroplasts with the MAR1-BINDING FILAMENT PROTEIN 1 (*At*MFP1) that is exclusively bound to thylakoid membranes (Seung *et al*., 2018). In *Atmfp1* and *Atptst2* mutants, most chloroplasts contain only a single, large starch granule, but this retains the WT-like lenticular shape and smooth surfaces (Seung *et al*., 2018; Thieme *et al*., 2022).

We investigated whether the effects of loss of *At*FZL on starch granule morphology result from its interactions with or displacement of the MFP1 thylakoid-binding component of granule initiation complexes. Our results provided no evidence for a functional link between *At*FZL and *At*MFP1. Although we were able to confirm that both *At*MFP1 and *At*FZL localise to puncta within Arabidopsis chloroplasts (Gao et al., 2006; Seung et al., 2018), co-expression of AtFZL-YFP and *At*MFP1-RFP in *N. benthamiana* chloroplasts revealed distinct punctate patterns for each protein, and the puncta overlapped only occasionally (Fig. 6). In addition, we did not detect a direct interaction of *At*FZL with AtMFP1 (Fig. S4). Moreover, *At*MFP1 localisation was still punctate in the *Atfzl* mutants (Fig. 6). These results support the idea that the defective starch granule morphology in *Atfzl* chloroplasts is caused by altered chloroplast architecture, rather than by direct interaction with *At*MFP1.

### Starch granule number in the Atfzl mutants correlates with chloroplast area

In common with plastid division mutants (Gao et al., 2006; Maple et al., 2007; Pyke et al., 1994; Shimada et al., 2004; Vitha et al., 2003), mesophyll cells of the *Atfzl-1* and *Atfzl-3* mutants contained larger chloroplasts with a broad variety of shapes. This resulted in the increase of the total chloroplast area per cell, when compared to the WT (Fig. 2). However, the size of the mesophyll cells was also increased in the *Atfzl* mutants (Fig. S2), leading to a similar chloroplast compartment size per cell as the WT. Interestingly, it seems that loss of *At*FZL does not affect the chloroplast size of epidermal pavement cells (Fig. 6).

Although the *Atfzl* mutants had a significantly broader distribution of starch granule numbers per chloroplast section than the WT (Fig. 2), the number of starch granules per unit chloroplast area in the mutants was not different from WT (Fig. 2). This result mirrors that of a previous study of Arabidopsis *arc* mutants defective in plastid division, where chloroplasts had approximately the same number of granules per unit volume of stroma as the wild type regardless of the size and number of chloroplasts per cell (Crumpton-Taylor *et al*., 2012). Thus, *At*FZL itself is not involved in the control of the number of starch granules per plastid volume. Likewise, the increased chloroplast size by itself is unlikely to cause the smaller starch granule size in the *A*t*fzl* mutants, since starch granule size is unaltered in the plastid division mutants *Atarc3* and *Atarc6* - which have enlarged chloroplasts comparable in size (or larger) to those of *Atfzl* mutants and their thylakoid membrane organisation remains largely unchanged (Maple *et al*., 2007; Robertson *et al*., 1995; Shimada *et al*., 2004; Vitha *et al*., 2003). More likely, the amount of accessible stromal volume or thylakoid membrane space might be involved in determining the number of initiation events per plastid. Future work using high-resolution 3D-imaging of mesophyll cells and chloroplasts in relation to the whole leaf will be necessary to further elucidate the relationships between cell and plastid size and starch granule formation.

### Leaf starch content is unaffected by the Atfzl mutations

The loss of proper thylakoid organisation in Arabidopsis can lead to a range of growth phenotypes – from severely retarded growth and development (e.g., mutants of VESICLE-INDUCING PROTEIN IN PLASTIDS 1-Gupta et al., 2021; Kroll et al., 2001) and variegated leaf patterns (e.g., mutants of THYLAKOID FORMATION 1 – Wang et al., 2004), to almost no defects in growth or leaf colour (CURVATURE THYLAKOID 1 – Armbruster et al., 2013). Arabidopsis *Atfzl* mutants were previously reported to have pale leaves, chlorotic lesions, reduced growth and delayed flowering (Gao et al., 2006; Landoni et al., 2013). However, under our growth conditions, they showed no differences in growth or development from the WT (Fig. 1 b-c). One major difference in growth conditions between our study and previous studies is that we used a 12 h rather than a 16 h photoperiod. The total starch content of *Atfzl* rosettes in our study was indistinguishable from the WT at the end of the day (Fig. 1 d-e). The absence of differences in both rosette fresh weight and starch content indicates that the *Atfzl* mutation is unlikely to have disrupted general photosynthetic efficiency and carbon fixation under our growth conditions. Interestingly, we observed a nearly complete degradation of starch granules in the *Atfzl* mutants during the night, whereas in the WT, small granules remained. It is possible that the reduced starch granule size allows more complete starch degradation in the mutants (Fig. 1e, 3). The opposite trend has been observed in mutants with increased granule size, such as *Atptst2, Atmrc* and *Atmfp1*, which all have significantly higher starch content than the WT at the end of the night (Seung *et al*., 2017; Seung *et al*., 2018).

In summary, we discovered that starch granules in Arabidopsis *Atfzl* mutants are smaller and have a more uneven surface morphology than the WT. While the experiments with *At*MFP1 seem to indicate that there is no direct association of *At*FZL with starch granule initiation proteins, our results suggest a strong influence of chloroplast ultrastructure and specifically thylakoid organisation on starch granule morphology.

## Materials and Methods

### Plant material and growth conditions

*Arabidopsis thaliana* plants were grown on soil in controlled environment chambers at constant 20°C and 60% relative humidity, and 12h light (80-150 µmol m^-2^ s^-1^)/12h dark cycles. *Nicotiana benthamiana* plants were grown in the glasshouse set to a minimum of 16 h light at 22°C.

Arabidopsis T-DNA insertion lines SALK_009051 (*Atfzl-1*), SALK_033745 (*Atfzl-2*) and SALK_152584C (*Atfzl-3*) were obtained from the Eurasian Arabidopsis Stock Centre (uNASC). Mutants SALK_009051, SALK_152584C were homozygous and mutant SALK_033745 was heterozygous as determined by genotyping PCR using the primers listed in Supplementary Table S1. Homozygous SALK_033745 plants were selected in the next generation by genotyping PCR using the primers listed in Supplementary Table S1.

### Cloning and plant transformation

To produce the *At*FZL-YFP construct, *At*FZL (without its stop codon) was amplified from total cDNA prepared from Arabidopsis leaves using the attB-flanked primers listed in Supplementary Table S1. The amplicon was recombined into the Gateway entry vector pDONR221 using BP clonase II (Thermo Fisher Scientific), and the sequence was verified by Sanger sequencing. The insert was recombined into the Gateway-compatible destination vector pUBC-YFP (Grefen *et al*., 2010) using Gateway LR clonase II (Invitrogen, Thermo Fisher Scientific). For *At*MFP1-YFP and *At*MFP-RFP constructs, we used the *At*MFP1 pDONR221 from Seung *et al*. (2018) to recombine the AtMFP1 coding sequence into pUBC-YFP and pB7RWG, respectively.

*Nicotiana benthamiana* leaves were transiently transformed via infiltration of *Agrobacterium tumefaciens* (GV3101) carrying the respective constructs. The bacteria were grown at 28 °C for 48 h. Cultures were resuspended in MMA buffer (10 mM MES pH 5.6, 10 mM MgCl_2_, 0.1 mM acetosyringone) at OD_600_ = 1.0 for infiltration of leaves to be used for confocal microscopy, and OD_600_= 0.3 (0.2 for p19) for leaves to be used for protein extraction. Leaves were infiltrated into the abaxial side using a syringe and harvested for confocal microscopy and protein extraction 48-72 h after infiltration.

*Arabidopsis thaliana* Col-0 and *Atfzl-3* mutant plants were transformed with *At*MFP1:pUBC-YFP by floral dipping (Zhang *et al*., 2006). Transformants were selected using the BASTA resistance marker.

### Starch content and iodine staining

Total starch content was quantified as described by Smith & Zeeman, 2006. Rosettes of 3-4 week old Arabidopsis plants were harvested at the end of day and end of night. The entire rosette was homogenised in 0.7 M perchloric acid and the insoluble fraction was collected by centrifugation. The pellet was washed in 80% ethanol three times and resuspended in water. Starch was gelatinised at 95°C and digested to glucose using a mix of amyloglucosidase (Megazyme) and ɑ-amylase (Megazyme). Glucose was then quantified by measuring NADH production using an assay based on hexokinase and glucose-6-phosphate dehydrogenase (Roche). Starch content was calculated in glucose equivalents.

For iodine staining, rosettes were harvested at the end of the day and chlorophyll was removed using 80% (v/v) ethanol prior to staining with Lugol’s solution (I_2_/KI solution).

### Chloroplast visualisation using light- and electron microscopy

Leaf segments were taken 1 mm from the central axis of Arabidopsis leaves of 3-4 week old plants at about one third of the distance between the tip and the petiole. Preparation for microscopy was as described in Watson-Lazowski et al. (2022). Briefly, leaf segments were fixed in 2.5% (v/v) glutaraldehyde in 0.05 M sodium cacodylate, pH 7.3 at 4°C, post-fixed in 1% (w/v) osmium tetroxide (OsO_4_) in 0.05 M sodium cacodylate for 2 h at room temperature, dehydrated in ethanol and infiltrated with LR White resin (Agar Scientific, Stansted, UK), using a EM TP embedding machine (Leica, Milton Keynes, UK). LR White blocks were polymerised at 60°C for 16 h. For light microscopy, semi-thin sections (ca. 0.5 µm) were prepared and stained with 1% (w/v) toluidine blue and the Periodic Acid Schiff kit (ab150680, Abcam) as described in Hawkins et al. (2021) and mounted in Histomount. Sections were imaged using the Zeiss Axio Imager Z2 using a 100x oil immersion objective. For transmission electron microscopy (TEM) ultrathin sections (ca. 80 nm) were cut with a diamond knife and placed onto formvar and carbon coated copper grids (EM Resolutions, Sheffield, UK). The sections were stained using 2% (w/v) uranyl acetate for 1 h and 1% (w/v) lead citrate for 1 min, washed in water and air dried. Sections were imaged on a Talos 200C TEM (FEI) at 200 kV and a OneView 4K x 4K camera (Gatan, Warrendale, PA, USA). Images were processed and analysed using ImageJ software (http://rsbweb.nih.gov/ij/) and Adobe Photoshop 2020. Granule numbers per plastid section were compared by fitting Mixed Effect Models (negative binomial regression) as in Chen et al. (2022).

### Starch granule purification, morphology and size analysis

Four-week-old plants (60 rosettes per genotype) were pooled and homogenised in 50 mM Tris-HCl, pH 8.0, 0.2 mM EDTA, 0.5% v/v Triton X-100 using an immersion blender. The suspension was sequentially filtered through Miracloth, a 60 µm nylon mesh and a 20 µm nylon mesh. Starch granules were separated by centrifugation at 2500*g* over a Percoll cushion (95% v/v Percoll, 5% v/v 0.5 M Tris-HCl, pH 8.0) then washed in water until clean, then washed twice in 0.5% w/v SDS in water and once in 100% ethanol to dry. Starch granule morphology was imaged using a Nova NanoSEM 450 (FEI) scanning electron microscope. Granule size distribution was analysed using a Beckman Multisizer 4e Coulter counter (Beckman Coulter) with a 30 µm aperture. A minimum of 50,000 particles were measured per replicate. Measurements were conducted with logarithmic bin spacing but are presented on a linear x-axis. Mean granule diameter was determined by the Multisizer 4e Coulter counter Software (Beckman Coulter).

### Confocal microscopy

For imaging of fluorescent proteins in Arabidopsis and *Nicotiana benthamiana* leaves, images were acquired on either the Zeiss LSM880 or the Leica Stellaris 8 laser scanning confocal microscope using a 40x or 63x water-immersion objective. YFP signal was excited using an Argon or a white light laser set to 514 nm and emission was detected at 519 nm to 560 nm. RFP signal was excited using a white light laser set to 555 nm and emission was detected at 562 nm to 623 nm. Chlorophyll autofluorescence was excited using 561 nm laser or a white light laser set to 555 nm, 561 nm or 576 nm and emission was detected at 651 nm to 750 nm. Images were processed using ImageJ software (http://rsbweb.nih.gov/ij/) and Adobe Photoshop 2020.

### Protein extraction and Immunoblotting

For the pairwise immunoprecipitation assay between *At*FZL-YFP and *At*MFP1-RFP, proteins were extracted from transiently transformed *Nicotiana benthamiana* leaves. One or two leaf discs (1 cm diameter) were harvested from each of three transformed leaves and homogenised in extraction buffer (50 mM Tris-HCl, pH 8.0, 150 mM NaCl, 1% v/v Triton X-100, 1x protease inhibitor cocktail (Roche), 1 mM DTT). The homogenate was centrifuged at 20,000*g* for 10 min, and the supernatant with the proteins was used as the input for immunoprecipitations. Immunoprecipitation was performed using the µMACs GFP Isolation Kit and µMacs Columns (Miltenyi Biotec). For immunoblotting, antibodies were used in the following concentrations: anti-YFP (Torrey Pines TP401 – 1:500) and anti-HA (Abcam ab9110 – 1:5000). Bands were detected using chemiluminescence, using the Anti-rabbit IgG (whole molecule)-Peroxidase (Sigma A0545 – 1:20000) and the SuperSignal West Femto Kit (Thermo Scientific).

## Acknowledgements

The authors thank the John Innes Centre (JIC) Horticultural Services for providing growth facilities and maintenance of plant material, JIC Bioimaging for providing access to microscopes, and Dr Andrew Breakspear for performing the LR-reaction to construct *At*MFP1 in pB7RWG and transforming it into *Agrobacterium*. We thank Prof. Alison Smith for critically reading this manuscript. This work was funded through a Leverhulme Trust Research Project grant RPG-2019-095 (to D.S), a John Innes Foundation (JIF) Chris J. Leaver Fellowship (to D.S), and BBSRC Institute Strategic Programme grants BBS/E/J/000PR9790 and BBS/E/J/000PR9799 (to the John Innes Centre).

## Author contributions

L.E. conceived the study. L.E. and D.S. designed the research. L.E., Q.Y.N. and J.E.B. performed the research and analysed data; L.E. and D.S. wrote the article with input from all authors.

## Figure Legends

**Figure S1: Starch granule number per chloroplast section in the *Atfzl-2* mutant**.

**(a-b)** Light microscopy images of *Atfzl-2* and wild type (Col-0) leaf sections. A young leaf was harvested from 29-day-old plants at the end of day. Sections were stained with toluidine blue and chi ‘s stain. Bar = 10 μm.

**(c-d)** Starch granule number per chloroplast section. Histograms show the frequency of chloroplasts containing a given number of granule sections relative to the total number of chloroplasts analysed (200 chloroplasts in one replicate).

**Figure S2: Starch granule number per area of chloroplast section and cell area**.

**(a-c)** Granule number per chloroplast plotted against chloroplast area for *Atfzl-1, Atfzl-3* and the wild type (Col-0). Starch granule number per chloroplast section (data in Figure 2d-f), and area of each chloroplast section was determined for 200 chloroplasts per biological replicate (I-III) per genotype.

**(d)** Area of cell section of 10 cells for each of three biological replicates. Values indicated with an asterisk are significantly different from Col-0 at each timepoint under a one-way ANOVA and Holm-Šídák multiple comparisons procedure (P ≤ 0.0).

**Figure S3: Chloroplast ultrastructure and starch granule placement in *Atfzl-2* mutant**.

TEM images of thin sections prepared from young leaves harvested from 29-day-old rosettes at the end o the day. rro heads indicate disor anised, ‘detached’ thylakoid layers o ser ed at the periphery of starch granules in *Atfzl-2*. Bar = 1 μm

**Figure S4: Pairwise immunoprecipitation of *At*FZL-YFP and *At*MFP1-RFP**.

Immunoprecipitation (IP) assay using anti-GFP beads for *At*FZL-YFP and *At*MFP1-RFP transiently co-expressed in tobacco leaves. Note that anti-GFP binds to *At*FZL-YFP but not *At*MFP1-RFP. Immunoblots with GFP and RFP antibodies were used to detect the *At*FZL-YFP and *At*MFP1-RFP proteins.

