## Supplemental Figures and Table for "A*t*FZL is required for correct starch granule morphology in Arabidopsis chloroplasts"

**Supporting information**

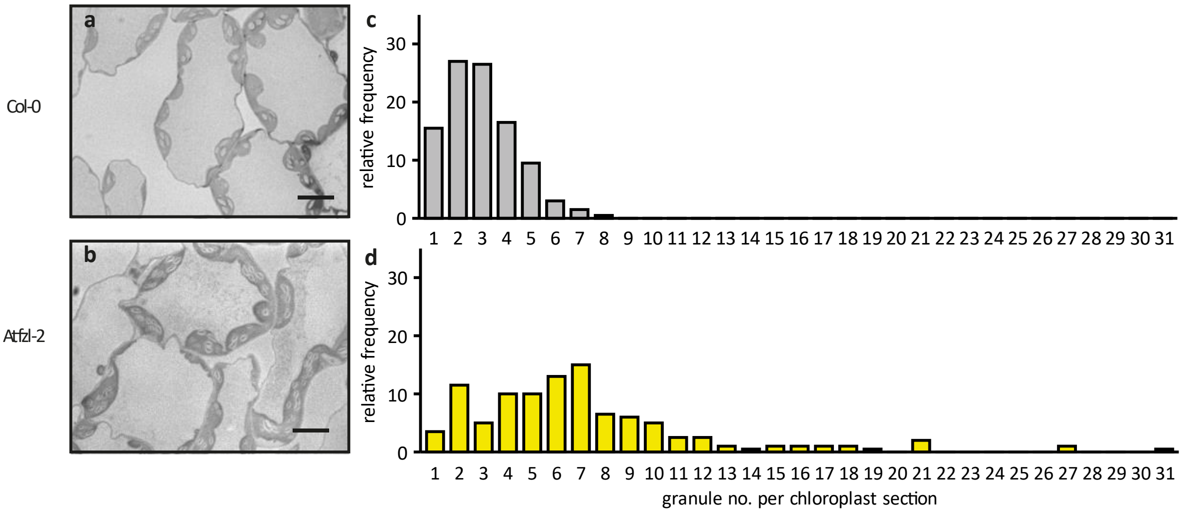


**Figure S1: Starch granule number per chloroplast section in the *Atfzl-2* mutant.**
**(a-b)** Light microscopy images of *Atfzl-2* and wild type (Col-0) leaf sections. A young leaf was harvested from 29-day-old plants at the end of day. Sections were stained with toluidine blue and Schiff's stain. Bar = 10 µm.
**(c-d)** Starch granule number per chloroplast section. Histograms show the frequency of chloroplasts containing a given number of granule sections relative to the total number of chloroplasts analysed (200 chloroplasts in one replicate).

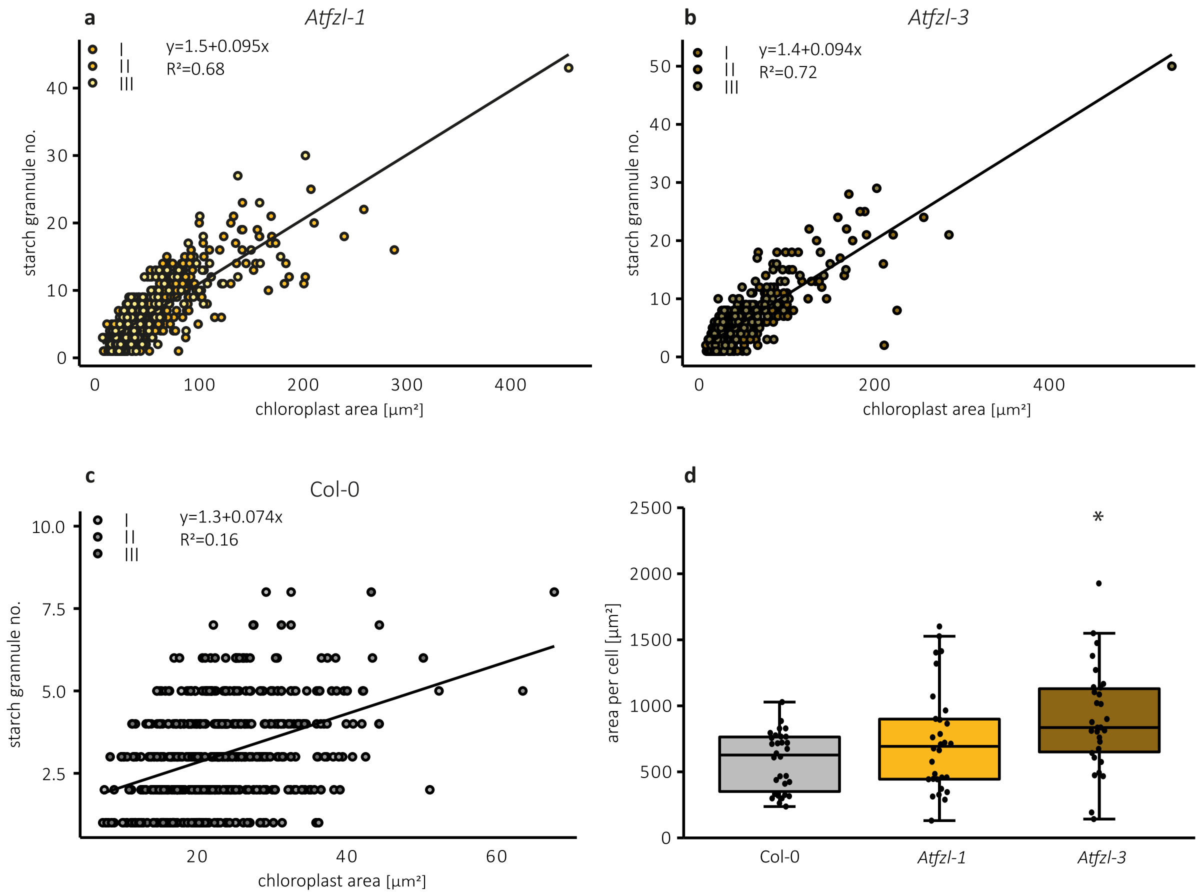


**Figure S2: Starch granule number per area of chloroplast section and cell area.**
**(a-c)** Granule number per chloroplast plotted against chloroplast area for *Atfzl-1*, *Atfzl-3* and the wild type (Col-0). Starch granule number per chloroplast section (data in Figure 2d-f), and area of each chloroplast section was determined for 200 chloroplasts per biological replicate (I-III) per genotype.
**(d)** Area of cell section of 10 cells for each of three biological replicates. Values indicated with an asterisk are significantly different from Col-0 at each timepoint under a one-way ANOVA and Holm-Šídák multiple comparisons procedure ($P \leq 0.05$).

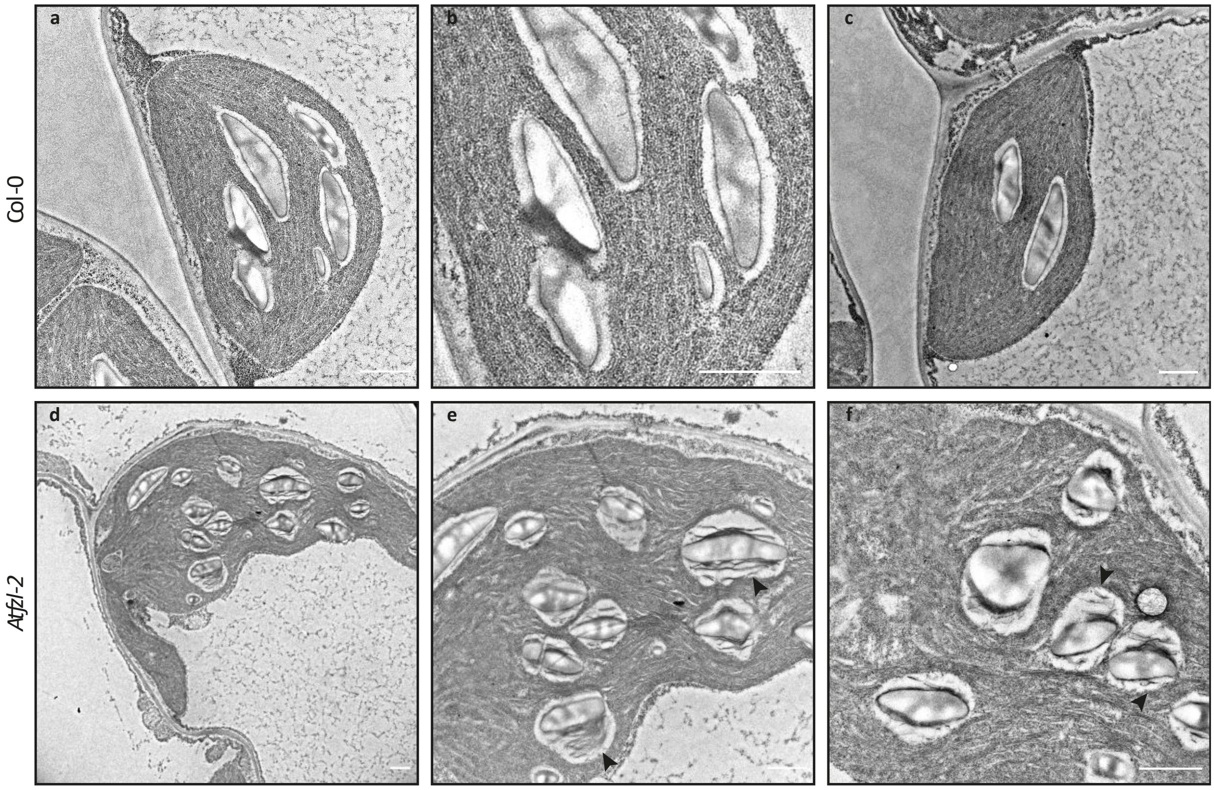


**Figure S3: Chloroplast ultrastructure and starch granule placement in *Atfzl-2* mutant.**
TEM images of thin sections prepared from young leaves harvested from 29-day-old rosettes at the end of the day. Arrowheads indicate disorganised, ‘detached’ thylakoid layers observed at the periphery of starch granules in *Atfzl-2*. Bar = 1 µm

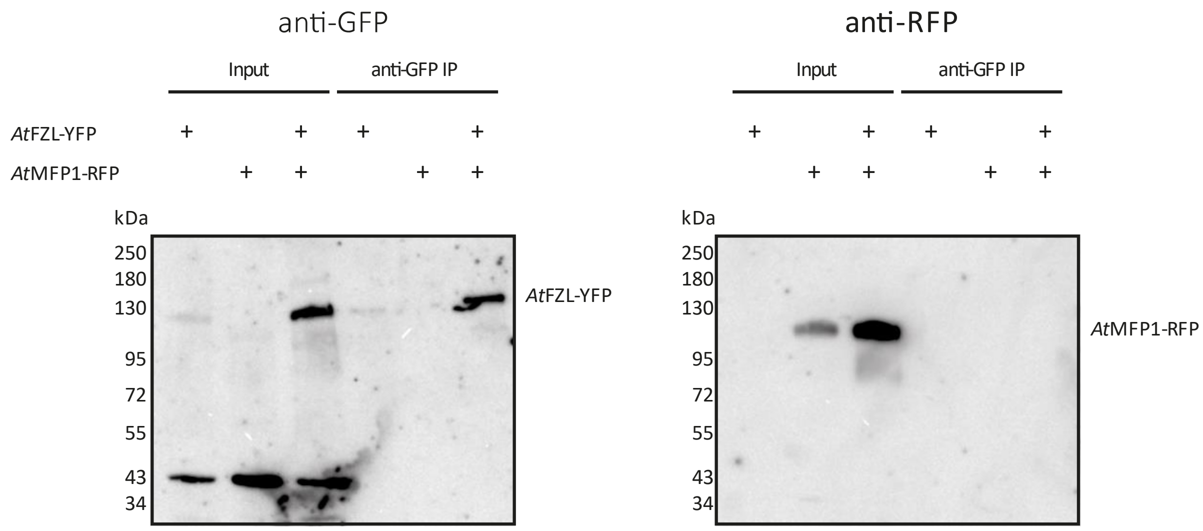


**Figure S4: Pairwise immunoprecipitation of *At*FZL-YFP and *At*MFP1-RFP.**
Immunoprecipitation (IP) assay using anti-GFP beads for *At*FZL-YFP and *At*MFP1-RFP transiently co-expressed in tobacco leaves. Note that anti-GFP binds to *At*FZL-YFP but not *At*MFP1-RFP. Immunoblots with GFP and RFP antibodies were used to detect the *At*FZL-YFP and *At*MFP1-RFP proteins.

**Table S1: Oligonucleotide primers for genotyping of *Atfzl* T-DNA mutants and cloning *AtFZL*.**

| genotyping | | |
|---|---|---|
| allele | forward primer | reverse primer |
| SALK_009051 | CCGAACCTATGATCTGACGG | AGCTTCCAACTCATCCAGC |
| SALK_033745 | CTCTTCCCAAGAAGTGCATTG | GATTCTGCTCTAATTGCCTCAA |
| SALK_152584C | ACTGCCGGAGAAAAAGAATTC | ATCTGCACGTGGAACAAATTC |
| Lba1 | TGGTTCACGTAGTGGGCCATCG | |
| cloning | | |
| gene | forward primer | reverse primer |
| AtFZL_At1g03160 | GGGGACAAGTTTGTACAAAAAAGCAGGCTTCACCATGAGAACTCTAATCTCTCACCGG | GGGGACCACTTTGTACAAGAAAGCTGGGTCAAGTCTCATCTCGTCTCGTGATA |

Lba1: T-DNA left border primer
